# DNA damage response clamp loader Rad24(Rad17) and Mec1(ATR) kinase have distinct functions in regulating meiotic crossovers

**DOI:** 10.1101/674507

**Authors:** Miki Shinohara, Douglas K. Bishop, Akira Shinohara

**Affiliations:** Institute for Protein Research, Osaka University, Osaka, 565-0871, JAPAN; School of Agriculture, Kindai University, Nara, 631-8505, JAPAN; Department of Radiation Oncology/Department of Molecular Genetics and Cell Biology, University of Chicago, Chicago, IL 60637, USA

**Author notes:** Correspondence: Akira Shinohara, Institute for Protein Research, Osaka University, 3-2 Yamadaoka, Suita, Osaka, 565-0871, Japan; Miki Shinohara School of Agriculture, Kindai University, 3327-204 Nakamachi, Nara, 631-8505, Japan.

**Keywords:** crossovers, crossover interference, Rad24, Mec1, meiosis

## Abstract

Crossover (CO) recombination is essential for chromosome segregation during meiosis I. The number and distribution of COs are tightly regulated during meiosis. CO control includes CO assurance and CO interference, which guarantee at least one CO per a bivalent and evenly-spaced CO distribution, respectively. Previous studies showed the role of DNA damage response (DDR) clamp and its loader in efficient formation of meiotic COs by promoting the recruitment of a pro-CO protein Zip3 and interhomolog recombination, and also by suppressing ectopic recombination. In this study, by classical tetrad analysis of *Saccharomyces cerevisiae*, we showed that a mutant defective in the *RAD24 gene* (*RAD17* in other organisms), which encodes the DDR clamp loader, displayed reduced CO frequencies on two shorter chromosomes (*III* and *V*) but not on a long chromosome (chromosome *VII*). The residual COs in the *rad24* mutant do not show interference. In contrast to the *rad24* mutant, mutants defective in the ATR kinase homolog Mec1/Esr1, including a *mec1* null and a *mec1* kinase-dead mutant, show little or no defect in CO frequency. On the other hand, *mec1* COs show defects in interference, similar to the *rad24* mutant. Moreover, CO formation and its control are implemented in a chromosome-specific manner, which may reflect a role for chromosome size in regulation.

## Introduction

Homologous recombination between two DNAs generates both crossover (CO) and non-crossover (NCO) products. CO recombinants display reciprocal exchange of the flanking DNA regions while NCO display uni-directional transfer of genetic information from one to the other parental DNAs with flanking regions remaining in their parental configuration. Meiotic recombination is regulated to promote the formation of COs while mitotic recombination is regulated to suppress them (Heyer *et al*. 2010; Hunter 2015). During meiosis COs play an essential role in correct segregation of homologous chromosomes during meiosis I (MI) for generation of gametes. As a consequence of their importance for chromosome segregation, the frequency and distribution of COs is strictly regulated during meiosis via a set of mechanisms referred to collectively as crossover control (Hunter 2015; Saito and Colaiacovo 2017).

Meiotic CO formation is initiated by the introduction of DNA double-strand breaks (DSBs) (Keeney 2001). Nucleolytic processing of DSB ends generates tracts of single-stranded DNA (ssDNA) that are substrates for the formation of nucleoprotein filaments by the strand exchange protein Dmc1 which then carries out a search for regions of homologous duplex DNA (Bishop *et al*. 1992). Invasion of the ssDNA into the homologous duplex results in the formation of an unstable displacement-loop (D-loop) (Hunter and Kleckner 2001). Then, a subset of D-loops is converted into a “stable” D-loop intermediates, that are destined to generate COs. Such CO intermediates are referred to as a single-strand invasion (SEI) (Hunter and Kleckner 2001). Further processing of SEIs produces double Holliday junction (dHJ) intermediates (Schwacha and Kleckner 1995). SEIs and dHJs are preferentially formed between homologous chromosomes rather than between sister chromatids (Schwacha and Kleckner 1994; Schwacha and Kleckner 1997). This is referred as to “interhomolog (IH) bias”. Finally, dHJs are biased to give rise to CO rather than NCO products (Allers and Lichten 2001; Hunter and Kleckner 2001). NCO recombinants are usually produced through the synthesis-dependent strand annealing pathway that does not form stable SEIs and dHJs (Allers and Lichten 2001; Hunter and Kleckner 2001; Borner *et al*. 2004; Mcmahill *et al*. 2007).

CO formation during meiotic prophase I is fostered by a group of proteins called ZMM (<u>Z</u>ip, <u>M</u>er, <u>M</u>sh) or SIC proteins (<u>S</u>ynaptic Initiation <u>C</u>omplex), hereafter ZMM for simplicity (Chua and Roeder 1998; Agarwal and Roeder 2000; Borner *et al*. 2004; Shinohara *et al*. 2008). ZMMs include Zip1, Zip2, Zip3, Mer3, Msh4, Msh5, Spo22/Zip4 and Spo16. Msh4-Msh5 and Zip2-Zip4-Spo16 sub-complexes bind to branched DNAs (Snowden *et al*. 2004; Arora and Corbett 2018; De Muyt *et al*. 2018). Mer3 encodes a 5’-3’ DNA helicase (Nakagawa and Ogawa 1999; Nakagawa *et al*. 2001). Zip3 is a putative SUMO/Ubiquitin E3 ligase (Agarwal and Roeder 2000; Perry *et al*. 2005; Cheng *et al*. 2006). ZMM proteins localize on meiotic chromosomes during prophase-I, which can be detected cytologically as immunostaining foci. Zip3, which is bound to DSB sites, promotes local assembly of other ZMM proteins (Shinohara *et al*. 2008; Serrentino *et al*. 2013). ZMM proteins also promote local initiation of synaptonemal complex (SC),(Sym *et al*. 1993; Storlazzi *et al*. 1996), and coordinate SC formation with recombination (Borner *et al*. 2004; Shinohara *et al*. 2008). ZMM proteins not only promote CO formation, but are also required for the normal distribution of meiotic COs (Sym and Roeder 1994; Chua and Roeder 1998; Agarwal and Roeder 2000; Novak *et al*. 2001; Shinohara *et al*. 2008).

Meiotic CO control involves three types of regulation. First, CO interference, which results in more even spacing of COs than expected at random (Muller 1916). Second, CO assurance distributes crossovers among chromosomes such that each chromosome receives at least one crossover, even when the average number of COs per chromosome in a single nucleus is low (Jones 1987; Bishop and Zickler 2004). Third, meiotic cells display “CO homeostasis”, a mechanism that buffers against low levels of dedicated CO intermediates by maintaining normal frequencies of COs at the expense of NCOs (Martini *et al*. 2006; Lao *et al*. 2013). Crossover frequency and control is also regulated in a chromosome-size (Kaback *et al*. 1992; Kaback *et al*. 1999) and also in per-nuclear basis (Wang *et al*. 2019). Cytological studies showed that the spacing of ZMM foci such as Zip3 indicates that crossover interference is established in *zmm* mutants, but that the sites designated to form COs are not maintained and many sites initially designated to become COs form NCOs instead, suggesting that ZMM is required for the maintenance of the class of COs that display CO interference (type 1 COs) (Fung *et al*. 2004; Zhang *et al*. 2014a). ZMM-dependent COs represent about 50-70% of the total number of COs observed in WT cells. The COs that remain in *zmm* mutants (type 2 COs) do not display interference. Importantly, topoisomerase II is necessary for interference between ZMM foci, indicating topoisomerase II is required to establish the regulated distribution of COs (Zhang *et al*. 2014b). CO interference is also regulated by proteins, which directly catalyze interhomolog recombination during meiosis (Shinohara *et al*. 2003b).

DNA damage response (DDR) proteins play an important role in the response to DNA damage in both mitotic and meiotic cells (Hochwagen and Amon 2006). During mitotic DDR in *S. cerevisiae*, the Rad24-RFC clamp-loader complex (the Rad17-RFC complex in other organisms) promotes recruitment of the Ddc1-Mec3-Rad17 clamp complex (“9-1-1” [Rad<u>9</u>-Rad<u>1</u>-Hus<u>1</u>] complex in other organisms) to tracts of ssDNA (Majka and Burgers 2003; Majka *et al*. 2006). Mec1(ATR) kinase is recruited on replication protein A (RPA)-coated ssDNAs through Ddc2(ATRIP) protein (Zou and Elledge 2003). Mec1 recruitment and activation are partly dependent on the clamp (Majka *et al*. 2006). In meiosis, as in mitosis, DDR proteins can induce delays in the entry into the first meiotic division (MI) when meiotic recombination is defective (Lydall *et al*. 1996; San-segundo and Roeder 2000). This delay during meiosis prophase I is imposed at the exit of pachytene, not at the onset of anaphase I (Subramanian and Hochwagen 2014). Cooperation of the DDR clamp and Mec1 activates a meiosis-specific CHK2 homolog, Mek1 kinase, which in turn relays signals to downstream targets (Hollingsworth and Gaglione 2019). One of Mek1 targets is Ndt80 transcriptional activator, which promotes pachytene exit by regulating the expression of genes required for late meiosis (Hollingsworth and Gaglione 2019). In addition to providing a means to regulate meiotic progression when recombination is incomplete, the DDR proteins work directly in meiotic recombination (Grushcow *et al*. 1999; Thompson and Stahl 1999; Shinohara *et al*. 2003a). The 911 clamp and Mec1 suppress non-allelic (ectopic) recombination (Grushcow *et al*. 1999; Thompson and Stahl 1999; Shinohara and Shinohara 2013). Furthermore, Mec1 promotes IH bias, presumably via its kinase activity, although the relevant substrates have yet to be identified, but given that Mec1 activates the down-stream kinase Mek1, and Mek1 is required for homolog bias (ref Kim/Kleckner-I think), it is likely that Mec1’s role in bias involves Mek1 activation.

Recently, we showed that 911 clamp, but not Mec1, promotes the loading of Zip3, thereby regulating ZMM assembly through direct protein interaction (Shinohara *et al*. 2015). Consistent with this, DNA analysis showed that the *911* and *rad24* mutants decreases the frequency of COs about 2-fold, as do *zmm* mutants (Grushcow *et al*. 1999; Shinohara *et al*. 2003a). Moreover, the *rad24* mutant is defective in IH bias (Shinohara *et al*. 2015). This particular feature of *rad24* is reminiscent of mutants lacking a functional *RAD51* gene; *RAD51* encodes a RecA homolog (Shinohara *et al*. 1992), that acts as an regulatory protein for the meiosis-specific RecA homolog Dmc1 during meiosis; Rad51 promotes Dmc1-mediated IH recombination (Schwacha and Kleckner 1997; Cloud *et al*. 2012; Hong *et al*. 2013). Over-expression of *RAD51* and its partner *RAD54* can suppress the reduced spore viability and X-ray sensitivity of *rad17* and *rad24* mutants (Shinohara *et al*. 2003a). Earlier studies in *Drosophila* showed that a mutant defective in the DDR kinase ATR homolog, *mei-41*, change the frequencies and distribution of meiotic COs along chromosomes (Carpenter and Baker 1982; Brady *et al*. 2018). However, the possible role of budding yeast DDR proteins in CO control has not been addressed.

In this paper, we provide genetic evidence for a novel role of the DDR clamp loader, Rad24, in efficient formation of meiotic COs on chromosomes *III* and *V*, which are relatively short chromosomes, but not on chromosome *VII*, one of the longest chromosomes. The COs that occur in the *rad24* mutant show little or no positive interference. Our results support the hypothesis that the clamp and clamp loader proteins promote interfering COs by promoting the recruitment of ZMM proteins to recombination sites. On the other hand, we find, although the *mec1* mutant displays reduced CO frequency on chromosome *III*, it displays increased CO frequency on chromosome *VII*. As with the *rad24* mutant, COs formed in the absence of Mec1 kinase activity display little if any interference. These observations suggest that Mec1 kinase regulates CO formation differentially from the DDR clamp loader. Mec1 may phosphorylate a novel protein, which is specifically involved in CO control, rather than being required for CO formation.

## Materials and Methods

### Strains and plasmids

All strains described here are derivatives of SK1 diploids except S2921 and MSY172, which are congenic strain (Sym and Roeder 1994). Strain genotypes are given in Table S1.

**Table 1.**
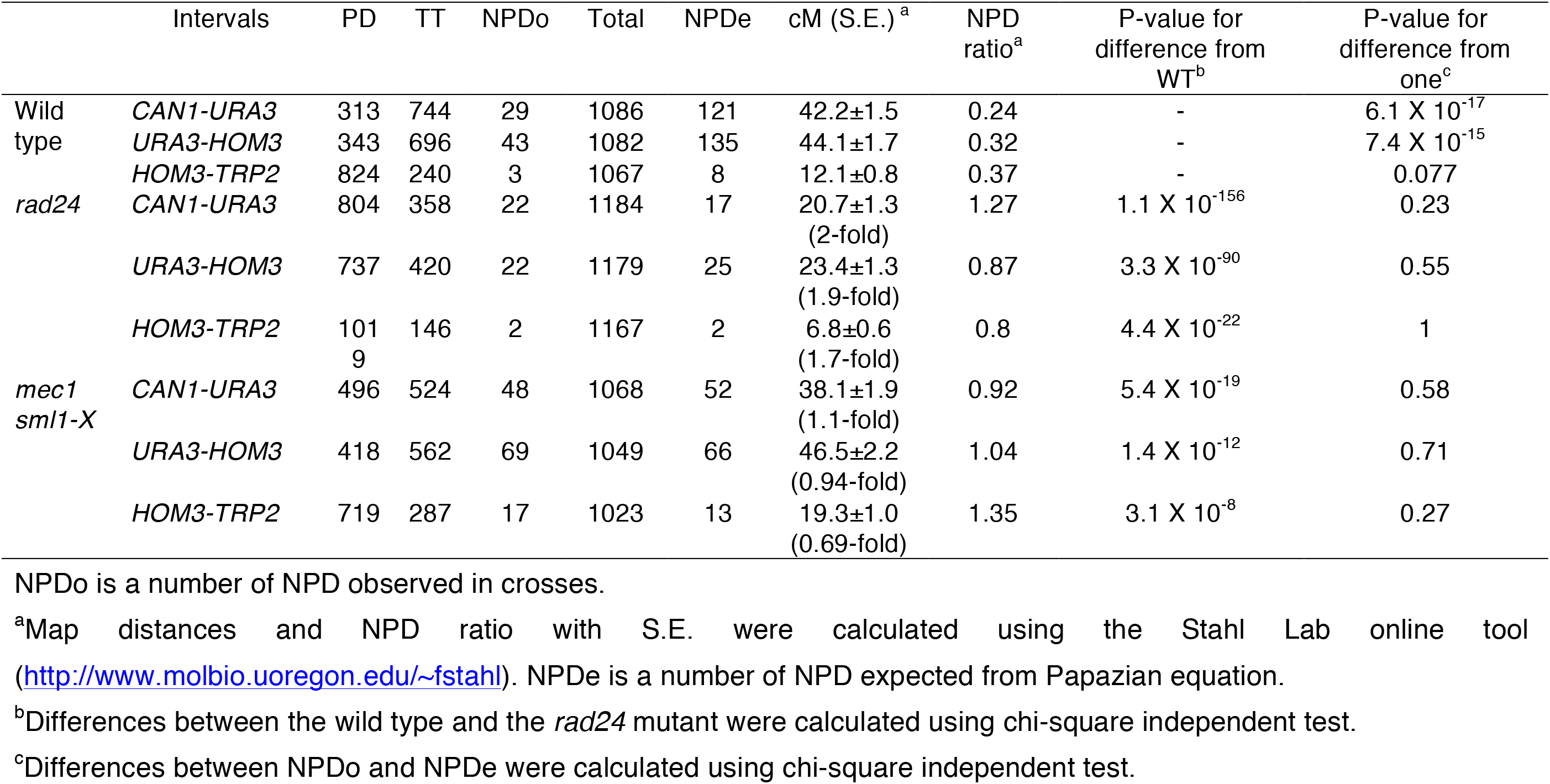
Genetic analysis of the meiotic recombination on chromosome *V* in *rad24* and *mec1* mutants

|  | Intervals | PD | TT | NPDo | Total | NPDe | cM (S.E.)<sup>a</sup> | NPD ratio<sup>a</sup> | P-value for difference from WT<sup>b</sup> | P-value for difference from one<sup>c</sup> |
| --- | --- | --- | --- | --- | --- | --- | --- | --- | --- | --- |
| Wild type | *CAN1-URA3* | 313 | 744 | 29 | 1086 | 121 | 42.2±1.5 | 0.24 | - | 6.1 X 10<sup>-17</sup> |
|  | *URA3-HOM3* | 343 | 696 | 43 | 1082 | 135 | 44.1±1.7 | 0.32 | - | 7.4 X 10<sup>-15</sup> |
|  | *HOM3-TRP2* | 824 | 240 | 3 | 1067 | 8 | 12.1±0.8 | 0.37 | - | 0.077 |
| *rad24* | *CAN1-URA3* | 804 | 358 | 22 | 1184 | 17 | 20.7±1.3<br>(2-fold) | 1.27 | 1.1 X 10<sup>-156</sup> | 0.23 |
|  | *URA3-HOM3* | 737 | 420 | 22 | 1179 | 25 | 23.4±1.3<br>(1.9-fold) | 0.87 | 3.3 X 10<sup>-90</sup> | 0.55 |
|  | *HOM3-TRP2* | 101<br>9 | 146 | 2 | 1167 | 2 | 6.8±0.6<br>(1.7-fold) | 0.8 | 4.4 X 10<sup>-22</sup> | 1 |
| *mec1*<br>*sml1-X* | *CAN1-URA3* | 496 | 524 | 48 | 1068 | 52 | 38.1±1.9<br>(1.1-fold) | 0.92 | 5.4 X 10<sup>-19</sup> | 0.58 |
|  | *URA3-HOM3* | 418 | 562 | 69 | 1049 | 66 | 46.5±2.2<br>(0.94-fold) | 1.04 | 1.4 X 10<sup>-12</sup> | 0.71 |
|  | *HOM3-TRP2* | 719 | 287 | 17 | 1023 | 13 | 19.3±1.0<br>(0.69-fold) | 1.35 | 3.1 X 10<sup>-8</sup> | 0.27 |

NPDo is a number of NPD observed in crosses.
<sup>a</sup>Map distances and NPD ratio with S.E. were calculated using the Stahl Lab online tool (http://www.molbio.uoregon.edu/~fstahl). NPDe is a number of NPD expected from Papazian equation.
<sup>b</sup>Differences between the wild type and the *rad24* mutant were calculated using chi-square independent test.
<sup>c</sup>Differences between NPDo and NPDe were calculated using chi-square independent test.

### Analyses of meiotic recombination

Time-course analyses of DNA events in meiosis were performed as described previously (Storlazzi *et al*. 1995; Shinohara *et al*. 1997). Genomic DNAs prepared were digested with *Mlu*I, *Bam*HI and *Xho*I (for heteroduplex) and *Pst*I (for meiotic DSB). Probes for Southern blotting were Probe “155” for HD and Probe 291 for DSB detection as described in (Storlazzi *et al*. 1995). Image gauge software (Fujifilm Co. Ltd., Japan) was used for quantification for bands of R1, R3 and DSB I.

### Genetic analysis of meiotic recombination

Genetic distances between markers and interference were analyzed as described before (Shinohara *et al*. 2003b; Shinohara *et al*. 2008).

Parental haploid strains were mated for 3 h on YPAD plates at 30°C and transferred onto SPM plates. After incubation at 30°C for 48 h, tetrads were dissected onto YPAD and incubated for 2 days. Genotyping was carried out as described (Shinohara *et al*. 2003a). In order to avoid aberrant clones (e.g. those containing mitotic crossovers), at least four independent crosses were carried out.

For interference analysis and genetic distance calculations, tetrads with non-Mendelian segregation of a diagnostic marker were excluded from the analysis. Map distances were determined using the standard mapping equation [cM]=100/2(TT+6NPD)/(PD+TT+NPD). Standard errors were calculated using the Stahl Lab online tool (http://www.molbio.uoregon.edu/~fstahl). Interference values are expressed as the NPD ratio; the fraction of tetrads expected to be NPDs was determined from the Papazian equation: NPDex=1/2[1-TT-(1-3T/2)^2/3^], where TT is the proportion of tetratypes observed (Papazian 1952). Data sets were analyzed using chi-square test.

To measure coincident double crossovers in adjacent intervals, the frequencies of tetrads with recombination in each of the two intervals were determined by summing TT and NPD tetrads for those intervals and dividing by the total number of tetrads (Shinohara *et al*. 2003b). The expected frequency of coincident recombination is given by the product of two single-interval frequencies.

### Statistical analysis

Graphs were prepared using and Microsoft Excel and GraphPad Prism 7. Datasets were compared using the Mann-Whitney U-test. χ^2^-test was used for proportion. Multiple test correction was done with Bonferroni’s correction. *, **, and *** show *P*-values of <0.05, <0.01 and <0.001, respectively.

### Data availability

Strains and plasmids are available upon request. The authors affirm that all data necessary for confirming the conclusions of the article are present within the article, figures, and tables

## Results

### The *rad24* mutant is defective in crossover formation and interference

Physical analyses have shown that the 9-1-1 clamp and its loader mutants in budding yeast *Saccharomyces cerevisiae* reduce crossover formation at a recombination hotspot on chromosome *III* (see below) (Grushcow *et al*. 1999; Shinohara *et al*. 2003a). To further characterize checkpoint mutant defects in crossover formation and control, we carried out tetrad analysis of meiotic recombination in the *rad24* mutant. For the analysis, we used markers on chromosome *V* in strains congenic with SK1 (Figure 1A) (Sym and Roeder 1994; Shinohara *et al*. 2003b). We dissected 1196 and 8060 tetrads for wild type and the *rad24* strains, respectively. Consistent with previous reports (Lydall *et al*. 1996; Shinohara *et al*. 2003a), the *rad24* mutant shows 32% spore viability and more tetrads having 0-, 2- and 4-viable spores per tetrad than those having 1- or 3-viable spores (Figure S1A); this pattern is consistent with loss of viability resulting from a high level of chromosome non-disjunction during the first meiotic division. Only about 1 in 6-7 tetrads formed by *rad24* mutants contained 4 viable spores. When genetic distances between markers on the chromosome are measured in 4-viable-spore tetrads, the *rad24* mutant reduces the distance of three intervals to about half of those in wild type (Figure 1, B and C and Table 1). Frequencies of non-Mendelian segregation for all four loci were also decreased 1.3 to 16-fold in the mutant (Table 2). The ability of a CO to interfere with coincident COs was determined by analysis of the ratio of observed to expected non-parental di-type (NPD) tetrads using the method of Papazian (see Materials and Methods). Wild type shows an NPD_observed_/NPD_expected_ (hereafter NPD) ratio of 0.37, 0.32, and 0.24 for the *CAN1-URA3*, *URA3-HOM3* and *HOM3-TRP2* intervals, respectively, indicative of CO interference. The *rad24* mutant exhibits an NPD ratio near unity for three intervals (1.27 0.87 and 0.8; Figure 2A and Table 1). This is the expected result for mutants defective in forming COs that display interference. These data show a role for Rad24 in controlling the distribution of COs along chromosomes. The defects in crossover formation and control in the checkpoint mutant are similar to those in the *zmm* core mutants such as *zip1* (Sym and Roeder 1994).

**Figure 1.**
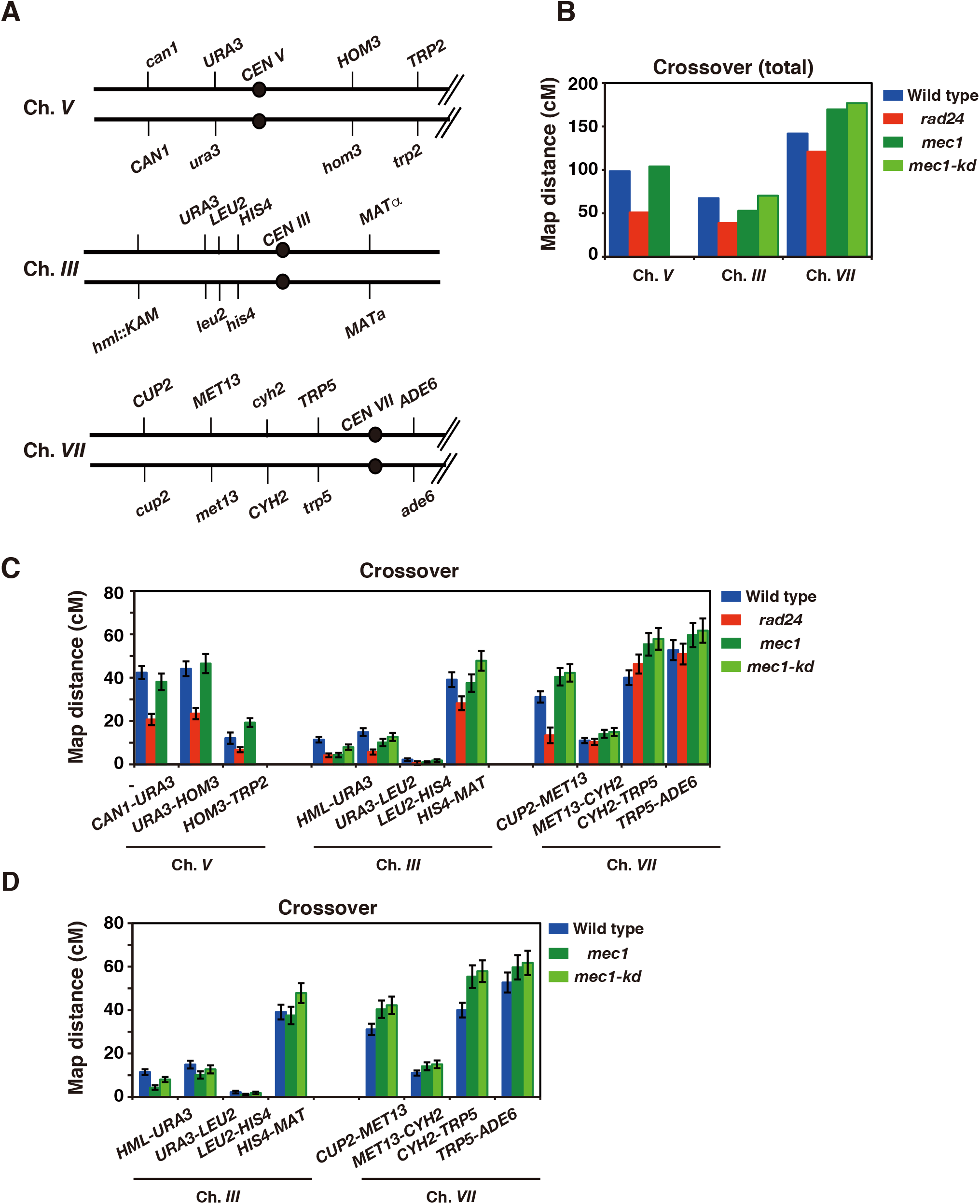
The *rad24* and *mec1* mutants are differentially defective in crossover frequency. (A) Schematic representation of markers on Chromosome *V*.
(B) Genetic maps of three different chromosomes in wild type (blue bars), the *rad24* (magenta bars), *mec1 sml1-X* mutants (green bars) and *mec1 sml1-X* mutants (light green bars). See Table 1, 3 and 4 for more details. Map distances were calculated using the Perkins equation.
(C) Genetic maps of different intervals on three different chromosomes in wild type (blue bars), the *rad24* (magenta bars), *mec1 sml1-X* mutants (green bars) and *mec1 sml1-X* mutants (light green bars). See Table 1, 3 and 4 for more details. Error bars indicate SE.

**Figure 2.**
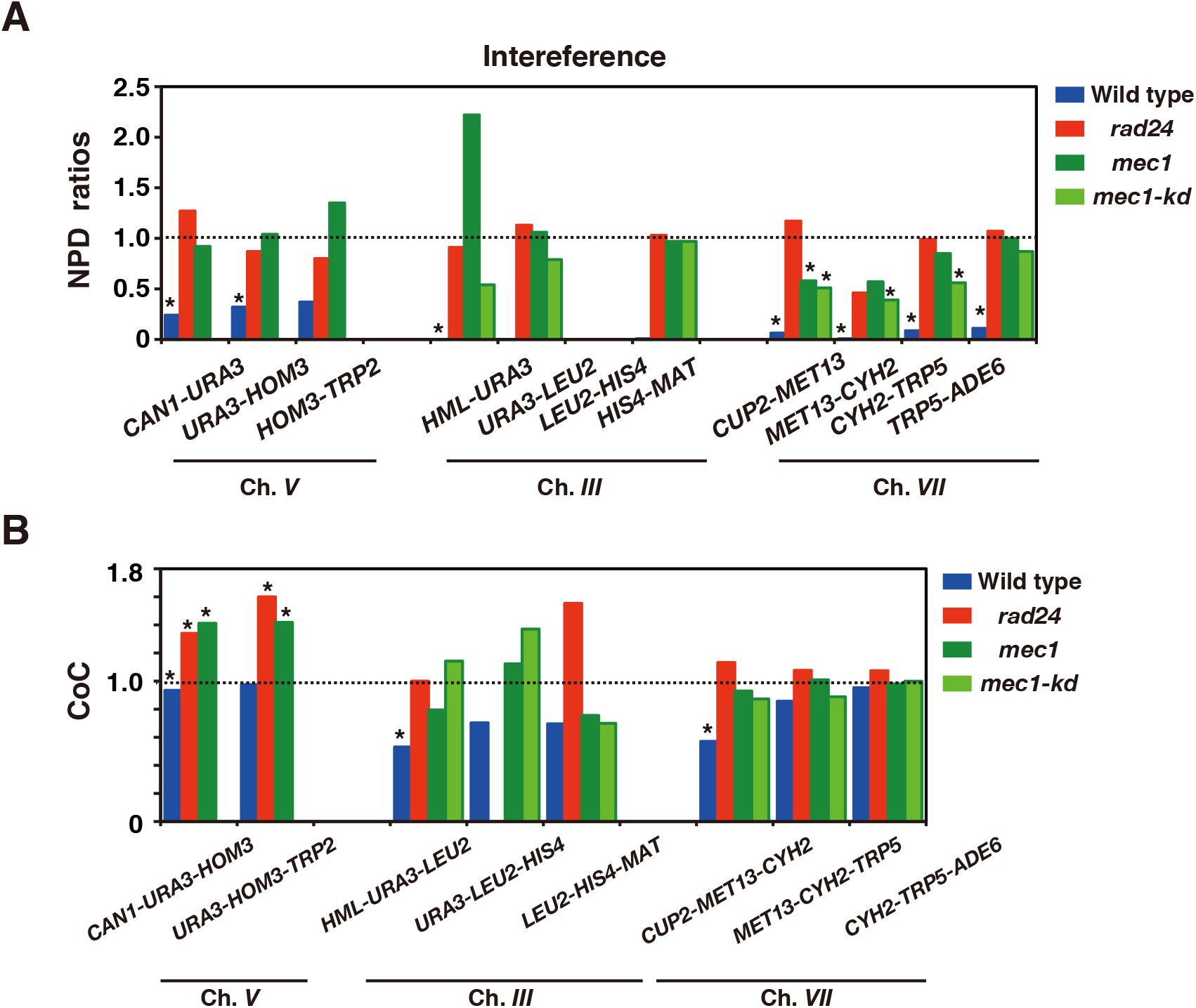
The *rad24* and *mec1* mutants are defective in crossover interference. (A) The NPD_observed_/NPD_expected_ ratios for eight intervals in wild type (blue), the *rad24* (red) a *mec1 sml1-X* mutants (green) and *mec1-kd sml1-X* (light green) were calculated as described in Methods and shown on chromosome *V*, *III* and *VII* (B). Also see Table 1, 3 and 4.
(B) Coefficient of coincidence (CoC) of CO frequencies of two adjacent intervals are calculated and shown on chromosome *V*, *III* and *VII*. Also see Table S2, S3 and S4.

**Table 2.** Non-Mendelian segregation in *rad24* and *mec1* mutants

|  | Percent of non-Mendelian segregation |  |  |  |
| --- | --- | --- | --- | --- |
|  | *CAN1* | *URA3* | *HOM3* | *TRP2* |
| Wild type | 1.2 | 0.3 | 1.5 | 1.8 |
| *rad24* | 0.67<br>(0.56) | 0.08<br>(0.27) | 1.17<br>(0.78) | 0.11<br>(0.06) |
| *mec1 sml1-X* | 1.2<br>(1) | 0.64<br>(2.1) | 2.9<br>(1.9) | 3.2<br>(1.8) |

Frequencies of tetrad with 4<sup>+</sup>:0<sup>-</sup>, 3<sup>+</sup>:1<sup>-</sup>, 1<sup>+</sup>:3<sup>-</sup> and 0<sup>+</sup>:4<sup>-</sup> segregation for each marker were calculated. Relative ratios of the frequencies in the mutant to wild type are shown in parenthesis.

NPD ratios detect interference between 4-strand double COs in a single interval, but they do not report on the interference that occurs between COs in two different intervals. The coefficient of coincidence [CoC] measures interference between COs in adjacent intervals. Interestingly, when the above tetrad data were analyzed for CoC, we noticed an unusual phenotype in the *rad24* mutant. Because two adjacent intervals used in this study are relatively long, wild type does not show any CO interference between two adjacent intervals (CoC is 0.90 and 0.98 for the *CAN1-URA3+URA3-HOM3* and *URA3-HOM3+HOM3-TRP2* interval pairs respectively; Figure 2B and Table S1). However, the *rad24* mutant displays negative interference; CoC in *CAN1-URA3-HOM3* and *URA3-HOM3-TRP2* is 1.33 and 1.6, respectively (Figure 2B and Table S2). These are significantly higher than 1 (*P*=8.0X10^-5^ for *CAN1-URA3+URA3-HOM3* and 1.5X10^-5^ for *URA3-HOM3+HOM3-TRP2*). Negative CO interference has been observed for another mutant strain (the *dmc1 hed1* double mutant)(Lao *et al*. 2013), which, like *rad24*, is defective in interhomolog partner choice. This finding suggests that *RAD24* may prevent clustering of COs on chromosome *V*.

### The *rad24* mutant is defective in CO interference on chromosomes *III* and ***VII***

We also analyzed several intervals on a short chromosome (Ch. *III*) and a long chromosome (Ch. *VII*) in a pure SK1 background (Higashide and Shinohara 2016) (Figure 1A). We dissected 1290 and 12090 tetrads for wild type and the *rad24* strains in this background, respectively. The *rad24* mutant showed reduced spore viability (31.2%; 97.7% in wild type; Figure S1B) in the pure SK1 background, which is similar to that in the congenic chromosome *V*-marked strain described above. The *rad24* mutant displayed CO frequencies in four intervals (*HML-URA3*, *URA3-LEU2*, *LEU2-HIS4*, and *HIS4-MAT*) that were roughly half as frequent overall as those in wild type (0.27-0.72 fold reduced; Figure 1, B and C, and Table 3). CO interference, as measured by NPD analysis, was almost compromised in the *rad24* mutant (Figure 2A and Table 3). This result is completely consistent with the corresponding data from the analysis of chromosome *V* in the congenic strain. When the intervals on chromosome *VII* were analyzed, the *rad24* mutant displayed a CO frequency in the *CUP2-MET12* interval that was roughly half that in wild-type (Figure 1C and Table 4). On the other hand, the other three intervals; *MET13-URA3*, *URA3-CYH2*, and *CYH2-ADE6* showed similar CO frequencies in wild-type and *rad24*. NPD ratios in three of the four intervals examined (Figure 2A and Table 4); *CUP2-MET12, URA3-CYH2* and *CYH2-ADE6* are almost one in the *rad24* mutant, indicating the absence of CO interference. Interference in the *MET13-URA3* was weakened relative to wild type (0.46 versus 0.063), but the data imply significant residual interference. These data suggest a chromosome (or an interval)-specific effect of the *rad24* mutation on CO frequency (see Discussion). We therefore determined CoC for three pairs of adjacent intervals on each chromosome *III* and *VII* in wild type (Figure 2B, and Table S3 and S4). With the exception of the *CYH2-TRP5+TRP5-ADE6* interval pair, five pairs showed CoC with less than one (Figure 2B), supporting the presence of CO interference between separated loci in wild type. In the *rad24*, the CoC values from four interval pairs are almost one or more than one. This supports the idea that COs formed in a given interval do not interfere with formation of COs in an adjacent interval in the absence of Rad24.

**Table 3.** Genetic analysis of meiotic recombination on chromosome *III* in *rad24* and *mec1* mutants

|  | Intervals | PD | TT | NPD<br>o | Total | NPDe | cM (S.E.)<sup>a</sup> | NPD ratio<sup>a</sup> | P-value for<br>difference from<br>WT<sup>b</sup> | P-value for<br>difference from<br>NPDe<sup>c</sup> |
| --- | --- | --- | --- | --- | --- | --- | --- | --- | --- | --- |
| Wild<br>type | *HML-URA3* | 1031 | 286 | 2 | 1301 | 9.30 | 11.4±0.65 | 0.0071 | - | 0.017 |
|  | *URA3-LEU2* | 908 | 312 | 7 | 1227 | 12.1 | 14.9±0.9 | 0.0099 | - | 0.14 |
|  | *LEU2-HIS4* | 1173 | 55 | 0 | 1228 | 0.31 | 2.2±0.3 | NA | - | 0.58 |
|  | *HIS4-MAT* | 533 | 712 | 49 | 1294 | 45.8 | 39.1±1.7 | 0.0069 | - | 0.64 |
| *rad24* | *HML-URA3* | 1191 | 104 | 1 | 1296 | 1.1 | 4.2±0.44<br>(0.37) | 0.91 | 9.0 X 10<sup>-75</sup> | 0.92 |
|  | *URA3-LEU2* | 1069 | 125 | 2 | 1196 |  | 5.7±0.57<br>(0.38) | 1.13 | 1.9 X 10<sup>-69</sup> | 0.88 |
|  | *LEU2-HIS4* | 1182 | 150 | 0 | 1197 | 0.02 | 0.6±0.43<br>(0.27) | NA | 8.4 X 10<sup>-15</sup> | 0.89 |
|  | *HIS4-MAT* | 734 | 500 | 36 | 1270 | 34.9 | 28.2±1.6<br>(0.72) | 1.03 | 3.2 X 10<sup>-33</sup> | 0.85 |
| *mec1*<br>*sml1-X* | *HML-URA3* | 1095 | 90 | 2 | 1187 | 0.90 | 4.3±0.52<br>(0.38) | 2.22 | 3.2 X 10<sup>-94</sup> | 0.25 |
|  | *URA3-LEU2* | 896 | 190 | 5 | 1091 | 4.70 | 10.1±0.84<br>(0.68) | 1.06 | 6.0 X 10<sup>-18</sup> | 0.89 |
|  | *LEU2-HIS4* | 1084 | 22 | 0 | 1106 | 0 | 1±0.21<br>(0.45) | NA | 4.8 X 10<sup>-14</sup> | NA |
|  | *HIS4-MAT* | 551 | 559 | 52 | 1162 | 53.8 | 37.5±2.0<br>(0.95) | 0.97 | 5.5 X 10<sup>-10</sup> | 0.81 |
| *mec1-kd*<br>*sml1-X* | *HML-URA3* | 1077 | 177 | 2 | 1186 | 3.68 | 8.0±0.6<br>(0.70) | 0.54 | 9.9 X 10<sup>-16</sup> | 0.38 |
|  | *URA3-LEU2* | 826 | 234 | 6 | 1066 | 7.59 | 12.7±0.94<br>(0.85) | 0.79 | 3.6 X 10<sup>-8</sup> | 0.56 |
|  | *LEU2-HIS4* | 1043 | 40 | 0 | 1083 | 0 | 1.8±0.29<br>(0.82) | NA | 3.0 X 10<sup>-6</sup> | NA |
|  | *HIS4-MAT* | 441 | 639 | 78 | 1158 | 80.2 | 47.8±2.3<br>(1.22) | 0.97 | 4.8 X 10<sup>-9</sup> | 0.81 |

NPDo is a number of NPD observed in crosses.
<sup>a</sup>Map distances and NPD ratio with S.E. were calculated using the Stahl Lab online tool (http://www.molbio.uoregon.edu/~fstahl). NPDe is a number of NPD expected from Papazian equation. The ratio of mutant to wild type map distance is shown in parenthesis.
<sup>b</sup>Differences between the wild type and the *rad24* mutant were calculated using chi-square independent test.
<sup>c</sup>Differences between NPDo and NPDe were calculated using chi-square independent test.

**Table 4.**
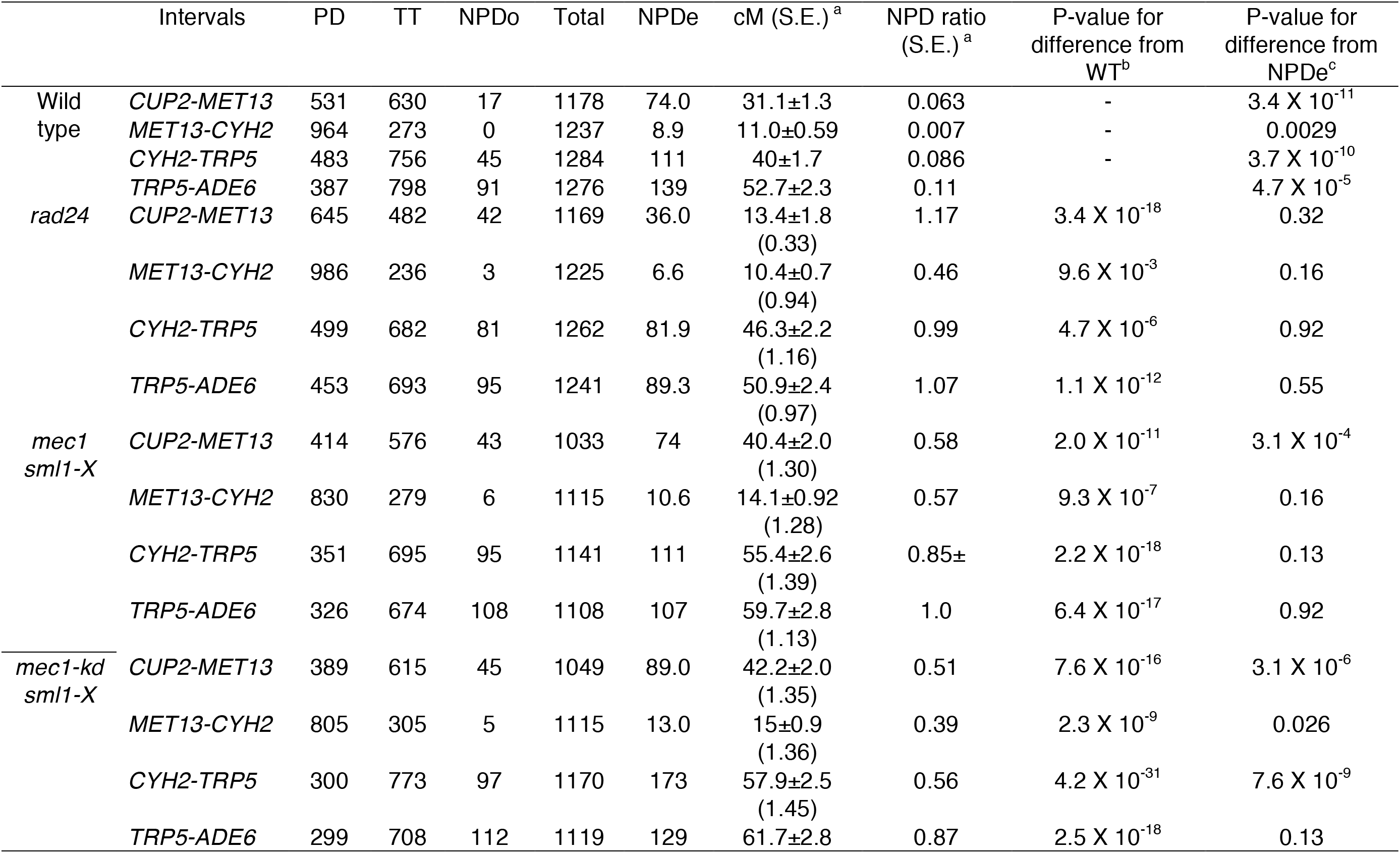
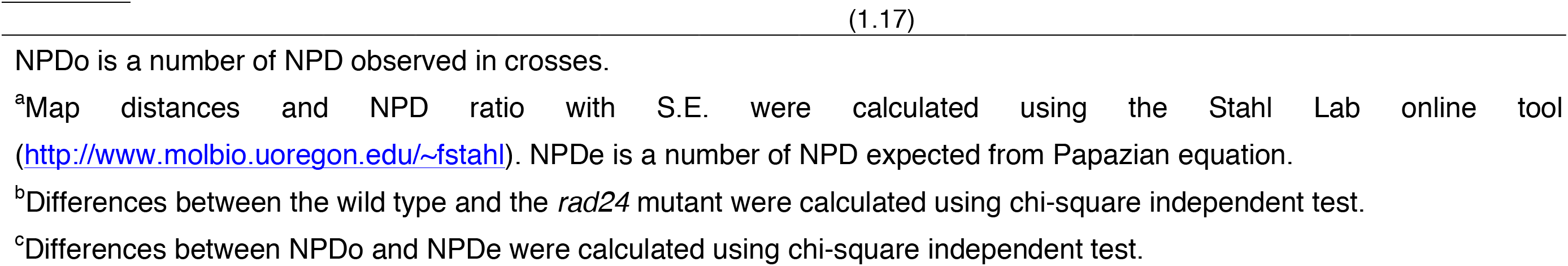
Genetic analysis of meiotic recombination on chromosome *VII* in *rad24* and *mec1* mutants

The *rad24* mutant displayed non-Mendelian segregation frequencies similar to wild-type at four of eight loci examined (Table 5). The other four loci showed an increase in the frequency of non-Mendelian segregation in the *rad24* mutant. These data indicate that the *rad24* mutant is proficient in NCO formation, but defective in CO formation on chromosome *III* and *VII*.

**Table 5.** Non-Mendelian segregation in *rad24* and *mec1* mutants

|  | Percent of non-Mendelian segregation<br>On Chromosome III |  |  |  |  | Percent of non-Mendelian segregation<br>On Chromosome VII |  |  |  |  |
| --- | --- | --- | --- | --- | --- | --- | --- | --- | --- | --- |
|  | *HML* | *URA3* | *LEU2* | *HIS4* | *MAT* | *CUP2* | *MET13* | *CYH2* | *TRP5* | *ADE6* |
| Wild type | 0.46 | 0.99 | 5.85 | 1.06 | 0.84 | 5.92 | 5.39 | 1.21 | 1.51 | 1.82 |
| *rad24* | 0.45 | 1.36 | 8.11 | 2.12 | 1.59 | 5.53 | 6.14 | 1.06 | 3.26 | 2.80 |
|  | (0.98) | (1.37) | (1.39) | (2.0) | (1.89) | (0.93) | (1.14) | (0.88) | (2.16) | (1.54) |
| *mec1* | 0.16 | 2.06 | 8.24 | 1.81 | 2.39 | 9.15 | 6.51 | 1.98 | 4.29 | 4.86 |
| *sml1-X* | (0.35) | (2.08) | (1.41) | (1.71) | (2.84) | (1.54) | (1.21) | (1.64) | (2.84) | (2.67) |
| *mec1-kd* | 0.08 | 3.18 | 10.4 | 2.20 | 3.59 | 7.17 | 7.91 | 1.55 | 3.18 | 5.71 |
| *sml1-X* | (0.17) | (3.21) | (1.78) | (2.08) | (4.27) | (1.21) | (1.46) | (1.28) | (2.11) | (3.14) |

Frequencies of tetrads with 4<sup>+</sup>:0<sup>-</sup>, 3<sup>+</sup>:1<sup>-</sup>, 1<sup>+</sup>:3<sup>-</sup> and 0<sup>+</sup>:4<sup>-</sup> segregation for each marker were calculated. Ratios of the frequencies in the mutant to wild type are shown in parenthesis.

### The *mec1* null mutant is defective in CO interference, but proficient in CO formation on chromosome *V*

Our previous study involving immunostaining of spread chromosomes revealed a difference between the effects of *rad24* and *mec1* mutations on chromosome-associated ZMM complexes, with only *rad24* conferring a defect in ZMM focus formation (Shinohara *et al*. 2015). We therefore analyzed *mec1* mutants in the congenic strain background for CO formation and control on chromosome *V*. The *mec1* deletion mutant was examined in combination with a *sml1-x* mutation, which suppresses the lethality conferred by the *mec1* by increasing levels of ribonucleotides (Zhao *et al*. 1998). The *mec1 sml1-x* double mutant shows 48.0% spore viability (5570 tetrads dissected), which is significantly higher than seen in the *rad24* mutant, consistent with a previous report (Lydall *et al*. 1996). Like the *rad24* mutant, the *mec1* deletion mutant produces more 4-, 2- and 0-viabile spore asci compared to the number of 3- and 1-viable spore asci (Figure S1A). Surprisingly, in obvious contrast to the *rad24* mutant, the *mec1* mutant displays the same map distance as wild type for one interval on Chromosome *V*, and a greater than wild-type map distance for two intervals (Figure 1C and Table 1). Moreover, the NPD ratio in the *mec1* mutant is essentially unity in two intervals, and is higher than wild type in a third interval (Figure 2A and Table 1). Furthermore, in contrast to the *rad24* mutant, the *mec1* mutant displays higher levels of non-Mendelian segregation than wild type (Table 2). Similar to the results obtained with the *rad24* mutant, the *mec1* mutant displayed negative interference, with CoC values exceeding unity (Figure 2B and Table S2). These findings suggest that *MEC1*(*ATR*) is not required for formation of normal levels of COs, but rather that the COs formed in *mec1* are distributed abnormally. Thus, the *mec1* mutant differs from the *rad24* mutant with respect to the ability to form normal levels of meiotic COs, but is similar in that it displays defects in positive interference, as well as apparent negative interference of distant coincident COs (See Discussion).

### The *mec1* mutants show chromosome-specific effects on CO formation

We also examined the effect of the *MEC1* deletion on meiotic recombination on chromosomes *III* and *VII* in a pure SK1 background. To rescue the lethality of the *MEC1* deletion, we introduced the *sml1-x* mutation and analyzed COs in 4-viable spore tetrads of *mec1-1 sml1-x* (hereafter, *mec1*). The *mec1* mutant showed reduced spore viability (60.2% in 3827 tetrad dissection) in the pure SK1 background, which is higher than that in the congenic background. On chromosome *III*, the *mec1* mutant reduced CO frequencies relative to wild type in three intervals; the ratio of CO frequency of mutant to wild type was 0.38 for *HML-URA3*, 0.68 for *URA3-LEU2*, and 0.45 for *LEU2-HIS4* (Figure 1, C and D, and Table 3). On the other hand, CO frequency in the *HIS4-MAT* in the *mec1-1* mutant is similar to that in wild type (0.95 fold). These indicate that the *mec1-1* mutant displays significant defects in the frequency of COs on chromosome *III* (Figure 1B). The NPD ratio in two intervals, *URA3-LEU2* and *HIS4-MAT*, was not significantly different from one in the *mec1-1* mutant (Figure 2A and Table 3), consistent with the absence of CO interference. The NPD ratio observed in the *HML-URA3* interval was 2.2, indicating negative interference (It should be noted, however, that this data set contains only 2 NPDs).

On chromosome *VII*, all four intervals, *CUP2-MET13, MET13-URA3*, *URA3-CYH2* and *CYH2-ADE6* show significant increase of CO frequencies in the *mec1* mutant compared to wild type; CO frequency ratios were 1.13-1.39 compared to wild type (Figure 1, B-D and Table 4). NPD ratios in the three intervals in the *mec1-1* mutant were all less than one (0.58, 0.57 and 0.85 in *CUP2-MET13, MET13-URA3*, *URA3-CYH2*, respectively), but significantly greater than those in wild-type. The NPD ratio for the *CYH2-ADE6* interval is about one in the *mec1-1* mutant. These data indicate that Mec1 is partially required for CO interference on chromosome *VII*. With respect to CoC values (Figure 2B and Table S3), *mec1-1* shows the same CoC value for the *LEU2-HIS4* and *HIS4-MAT* intervals as wild type. The CoC values obtained for the other three interval pairs examined were significantly higher in *mec1-1* than those in wild-type, and were close to one. This again supports the role of Mec1 in normal distribution of COs along chromosomes.

For non-Mendelian segregation, the *mec1-1* mutant showed increased frequencies at 7 of 8 loci examined on chromosomes *III* and *VII* (Table 5). The exception was the *HML* locus, which was assayed via marker insertion; the *mec1* mutation did not show a significant effect on non-Mendelian segregation frequency at that locus.

### Mec1 kinase activity is required for CO interference

Given that Mec1 belongs to a PI3-kinase family (Majka *et al*. 2006), we wondered whether the kinase activity of Mec1 is required for CO control during meiosis. We introduced the *mec1-kd* (kinase-dead; D2224A) allele in the pure SK1 background (with *sml1-x*) with markers on chromosomes *III* and *VII*. The *mec1-kd* mutant showed reduced spore viability (75.5% in 3218 tetrad dissection), which is higher than that in the *mec1-1* and *mec1* null mutants, suggesting the *mec1-kd* mutation is a hypomorphic mutant. On chromosome *III*, the *mec1-kd* mutant reduced CO frequencies compared to the wild type in three intervals, with frequency ratios of 0.7 for *HML-URA3*, 0.85 for *URA3-LEU2*, and 0.82 for *LEU2-HIS4* (Figure 1, C and D, and Table 3). On the other hand, CO frequency in the *HIS4-MAT* in the *mec1-kd* mutant is slightly higher than that in wild type (1.2 fold). Compared to the *mec1-1* mutant, the *mec1-kd* mutant showed weaker defects in CO formation. NPD ratios of *HML-URA3* and *URA3-LEU2* in the *mec1-kd* were 0.54 and 0.79 respectively, significantly higher than those in wild type, but significantly lower than one (Figure 2A and Table 3). Furthermore, the ratios are significantly lower than those in the *mec1-1* mutant. On the other hand, NPD ratio of the *HIS4-MAT* interval in the mutant is 0.97, indicating a strong defect in interference. Thus, the *mec1-kd* mutant shows weaker defects in CO formation and interference than *mec1-1* mutant. This suggests that there is significant residual CO interference in the *mec1-kd* relative to *mec1-1.* This may reflect residual kinase activity or a kinase-independent role of Mec1 in CO interference.

For the intervals on chromosome *VII*, the *mec1-kd* mutant showed increases in the frequency of COs for all intervals (1.17-1.45 fold; Figure 1, C and D, and Table 4), which is similar to the results obtained in the *mec1* null mutant. The NPD ratios of all four intervals in the *mec1-kd* mutant ranged from 0.39 to 0.87 (Table 4). These are values are significantly higher than those in wild type, but lower than those in the null mutant. Thus, the *mec1-kd* mutant displays weaker defects in CO control than the *mec1* null mutant. With respect to CoC values (Figure 2B and Table S3 and S4), among five pairs, the *mec1-kd* shows the same CoC value for the *LEU2-HIS4 and HIS4-MAT* intervals as wild type. The CoC values of the *HML-URA3: URA3-LEU2* pair of intervals and the *URA3-LEU2: LEU2-HIS4* pair of intervals on chromosome *III* in the *mec1-kd* mutant are significantly higher than those in the wild-type, which are more than one, again indicating the absence of interference between CO events in adjacent intervals. On chromosome *VII*, CoC of for the C*UP2-MET13:MET13-CYH2* interval pair in the *mec1-kd* is significantly higher than that in wild type. On the other hand, CoC value of the *MET13-CYH2:CYH2-TRP5* intervals in the *mec1* mutant is significantly, though modestly, higher than that in wild type. These data are consistent with the above result that the *mec1-kd* mutant showed a weaker defect in CO interference than the *mec1-1* mutant.

The *mec1-kd* mutation increased the frequencies of non-Mendelian segregation at 7 out 8 loci examined on chromosomes *III* and *VII*, with the exception being the *HML* locus (Table 5). This trend is similar to that observed in the null mutant.

### *RAD24* functions upstream of *ZIP1* in meiotic recombination

The above genetic analysis showed a similarity between *rad24* and the *zmm* mutants in terms of CO defects. To determine the relationship *DDR* and *ZMM* genes, we carried out DNA analysis. Previous physical analysis showed that the *ZIP1* and the *RAD24* genes work in the same pathway to suppress non-allelic meiotic recombination (Shinohara and Shinohara 2013). We checked whether *ZMM* and *DDR* genes work in the same pathway of recombination by analyzing meiotic recombination products of the *rad24 zip1* double mutant at a DNA level. We examined levels of CO and NCO heteroduplexes (HDs), as well as levels of ectopic (EC) recombinants at the *HIS4-LEU2* recombination hot spot on a single blot (Figure 3, A and B). The HD assay was used in place of the conventional NCO/CO assay (Storlazzi *et al*. 1995) because the HD assay resolves bands specific for EC from all NCO/CO bands (Shinohara and Shinohara 2013). Comparison of two phenotypes of the *rad24 zip1* double mutant with those of the corresponding single mutants shows that *rad24* and *zip1* affect meiotic recombination on the same pathway (Figure 3B). First, the *rad24 zip1* double mutant shows elevated levels of EC recombination like *rad24* rather than reduced levels like *zip1* (Figure 3, B and C). This is consistent with our previous study of direct measurement of EC (Shinohara and Shinohara 2013). Second, and particularly important to the central conclusion of this study, CO-HD levels in *zip1* are similar to those in the *rad24 zip1* double mutant (Figure 3C). Third, the *rad24 zip1* double mutant shows a decrease in the NCO/CO-HD ratio relative to wild type like the *rad24* single mutant, rather than an increase in the ratio like the *zip1* single mutant. The ratio of NCO/CO-HD is about 1.3 in WT cells. This ratio is reduced to 0.6 in the *rad24* mutant and increased to 3.5 in *zip1*. The NCO/CO-HD ratio in the *rad24 zip1* double mutant is about 0.6, equivalent to the *rad24* single mutant. These observations imply that the COs that form in *rad24* mutants are *ZIP1*-independent and that Rad24 and Zip1 contribute to the same pathway for CO formation. Furthermore, the results indicate that the increase in NCO levels observed in the *zip1* mutant depends on Rad24, suggesting that Rad24 functions upstream of Zip1 to channel intermediates to become interhomolog COs or NCOs rather than EC recombinants.

**Figure 3.**
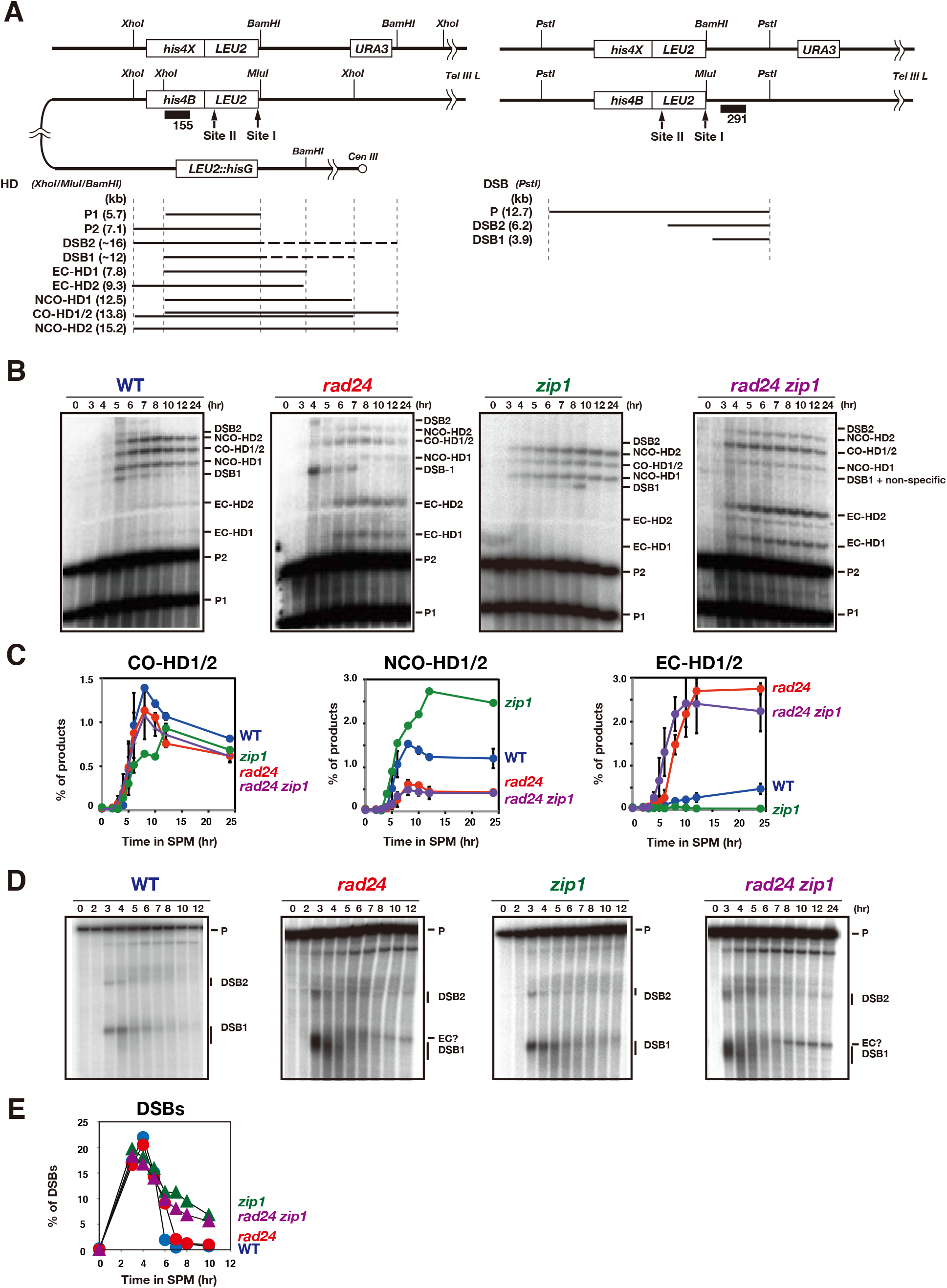
*RAD24* functions upstream of *ZIP1* in meiotic recombination. (A) Schematic representation of *HIS4-LEU2* and *leu2::hisG* loci. HD assay, left; DSB assay, right. In HD assay, DSB bands contain single-stranded DNA tails (shown by dotted lines; DSB1, DSB2), which confers resistance to digestion by restriction enzyme.
(B) Heteroduplex (HD) formation in wild type and the various mutants was analyzed by Southern blotting. Genomic DNAs were digested with *Bam*HI, *Mlu*I and *Xho*I.
(C) Quantification of NCO-, CO- and EC-HDs is shown in each graph. Wild type, blue (n=3); *rad24*, red (n=3); *zip1*, green (n=1); *rad24 zip1*, purple (n=3). Note: interpretation of HD band levels assumes that the mutations examined do not alter the efficiency of mismatch repair of the *Mlu*I/*Bam*HI HDs.
(D) DSB blots in various mutants. DSBs were analyzed by Southern blotting. Genomic DNAs were digested with *Pst*I. Variable resection of DNA ends is likely to account for the low levels of DSB-specific bands in *rad24 zip1*.
(E) Quantification of kinetics of DSB repair shown in (D). Wild type, blue circles: *zip1*, green triangles; *rad24*, red circles; *rad24 zip1*, purple triangles.

Finally, the *rad24 zip1* double mutant shows a very similar defect in DSB repair as the *rad24* single mutant. The double mutant also displayed similar hyper-resection compared to wild type and (the *zip1*) as the *rad24* single (Figure 3, D and E). These data lead us to conclude that the Rad24 clamp loader, and therefore probably the 9-1-1 clamp, promotes *ZIP1*-dependent CO events.

## Discussion

Rather than simply regulating cell cycle progression, DNA damage response (DDR) proteins play multiple roles in DNA metabolism during meiosis. These roles include suppression of non-allelic homologous recombination (NAHR) and DSB formation, as well as promotion of IH bias, CO formation, and CO control. The 9-1-1 clamp loader/clamp and Mec1(ATR) kinase have distinct functions during meiotic prophase I. Although both the 911 clamp loader/clamp and Mec1(ATR) are required for suppression of NAHR, normal levels of meiotic CO formation require DDR clamp loader/clamp, but not Mec1. The DDR clamp loader/clamp promotes the functions of ZMM proteins during CO formation, likely via its direct interaction with Zip3 (Shinohara *et al*. 2015). Although previous studies analyzed the role of DDR proteins in CO formation, the roles of DDR protein in CO control had not been tested. Here, we carried out genetic analysis of a *rad24* null mutant and various *mec1* mutants. Taken together, the data suggest that Rad24 and Mec1 play distinct roles in regulating meiotic CO formation and distribution. One type of CO control, CO interference, which can be measured genetically or cytologically, consists of at least two distinct stages; establishment and maintenance (Zhang *et al*. 2014a; Zhang *et al*. 2014b). Previous studies showed the role of ZMM proteins in the maintenance of the interference (Zhang *et al*. 2014a; Zhang *et al*. 2014b). In the *zmm* mutants, the sites of COs appear to be designated and display the even spacing indicative of interference, but are not matured into CO products, but rather into NCOs. Given the similarity in defects in CO formation and control between ZMM and DDR mutants, we speculate that, like ZMM, Rad24 and Mec1 are both involved in CO maintenance, but their roles in maintenance are distinct.

We carried out genetic analysis of viable spores of several DDR mutants after tetrad dissection. This method relies on spore viability. The reliance on 4-viable spore tetrads in mutants that reduce spore viability could have biased the results. On the other hand, we tested three different alleles of the *mec1* mutant, which show different spore viability from 48% of *mec1* null to 76% for *mec1-kd*. Importantly, as shown above, all three *mec1* mutants show the similar phenotype in terms of CO/NCO formation and CO interference. This argues against a strong impact of selecting for 4-viable spore tetrads on the recombination associated phenomena examined, at least in the case of the *mec1* mutants. Furthermore, selection for 4-viable-spore tetrads is likely to underestimate the impact of a recombination-defective mutant rather than exaggerate it.

The defect in CO interference in *rad24* is reminiscent of *zmm* mutants. On the other hand, our results reveal new differences between *rad24* and *zmm*. The *rad24* mutant reduced CO frequencies to half of those in wild type in all intervals on both chromosome *III* and *V*, but did not reduce CO frequencies in several intervals on a long chromosome, chromosome *VII*. COs on all three chromosomes in the *rad24* mutant do not show the interference. In contrast, a typical *zmm* mutant, *zip3*, reduced CO frequencies irrespective of chromosome size (Chen *et al*. 2008; Oke *et al*. 2014). One possible explanation for wild-type levels of COs on chromosome *VII* in the *rad24* mutant is that increases in ZMM-independent COs might compensate for loss of ZMM-dependent COs, but do so only on long chromosomes. A recent study showed that SC assembly down-regulates DSB formation during middle and late prophase I (Thacker *et al*. 2014). Given the *rad24* mutant is defective in SC formation (Grushcow *et al*. 1999), the mutant may display higher levels of DSB formation on longer chromosomes than shorter chromosomes in late prophase I due to incomplete synapsis. If so, such late DSBs might contribute to ZMM-independent CO formation, since the mutant is deficient in ZMM-dependent CO formation. In this scenario, Rad24 might block ZMM-independent COs that otherwise arise from late DSBs on long chromosomes. Alternatively, long chromosomes may have a property that allows recruitment of ZMM independently of Rad24.

The *rad24* and *mec1* mutations confer different defects with respect to CO frequency and distribution. The simple explanation on the *mec1* phenotypes in meiotic CO formation and control is that normal channeling of intermediates to the ZMM-dependent CO pathway, which is subject to interference, is lost in *mec1* such that COs occur by a ZMM-independent, non-interfering, CO pathway. The Mec1 may activate the ZMM pathway and/or suppress a ZMM-independent pathway. The *mec1* kinase dead mutant also shows CO phenotypes similar to those in the null mutant, suggesting the phosphorylation by Mec1 kinase is involved in regulation of COs. If Mec1 activates the ZMM-dependent pathway, such regulation is likely to occur after the assembly of ZMM complexes on chromosomes, because the *mec1* mutant is proficient in ZMM focus formation (Shinohara *et al*. 2015).

An alternative model is based on the observation that Mec1(ATR) suppresses DSB formation through DSB *trans*-interference (Zhang *et al*. 2011; Garcia *et al*. 2015). In this model, the *mec1* mutants undergo more DSB formation than wild type. Indeed, our genetic analysis indicates that *mec1* mutants display increased NCO frequencies at most loci assayed, which is a result expected if more DSBs occur per meiosis in a given interval. If the additional DSBs are processed via a ZMM-independent CO pathway, which also forms NCO, that would account for the *mec1* defects described in this study. Even in this scenario, Mec1 seems to select the fate of the additional DSBs that are resolved by either ZMM-dependent or ZMM-independent mechanisms.

A recent study of whole-genome recombination analysis using hybrid yeast strains reached the same conclusion as we do here regarding the distinct regulatory roles of *rad24* and *mec1* in CO formation and control during meiosis (Crawford…and Neale et al. BioRxiv).

The *mec1* mutants we examined display chromosome-specific defects in CO frequency, which may reflect a mechanism that depends on chromosome size. The data are consistent with Mec1 enhancing the frequency of CO formation on small chromosomes, such as chromosome *III*, while it suppresses CO formation on long chromosomes, such as chromosome *VII*. Consistent with this, the *mec1* mutant does not affect CO frequencies on a medium length chromosome, chromosome *V*. The different effects of Mec1 kinase on COs seen on different intervals on different chromosomes suggest Mec1 senses some chromosome- or interval-specific property, with regulation based on chromosome size likely. In this context it should be pointed out that previous studies by Kaback and colleagues showed that both CO frequency and the strength of interference depends on chromosome size (Kaback *et al*. 1992; Kaback *et al*. 1999). These studies showed that long chromosomes display more robust interference than small chromosomes. Our results are consistent with the possibility that Mec1 is involved in size-dependent control to distribute COs among different chromosomes such that shorter chromosomes are assured of undergoing the CO events required for their proper segregation. This type of control cannot be accounted for as a consequence of *trans* DSB interference, because DSB interference only acts over short distances on a given chromosome (Garcia *et al*. 2015).

In closing we emphasize that, although our data demonstrate distinct roles for Rad24 and Mec1 in generating the normal distribution of CO events, genetic data of the type presented here cannot distinguish between interference mutants that fail to establish a non-random pattern of potential CO sites and interference mutants that fail to maintain the non-random pattern until CO products are eventually formed (Zhang *et al*. 2014a). Cytological studies will be required to determine if *RAD24* and/or *MEC1* play a role in establishment verses maintenance of CO patterns.

## ACKNOWLEDGEMENTS

We acknowledge Dr. N. Hunter for critical reading and are grateful for Dr. Matt Neale for sharing unpublished results prior to publication. We thank Ms. A. Murakami and H. Wakabayashi for excellent technical assistance. This work was supported by JSPS KAKENHI Grant Number; 22125001, 22125002, 15H05973 and 16H04742 to A.S. M.S. was supported by the Japan Society for the Promotion of Science (JSPS) through the Funding Program for Next Generation World-Leading Researchers (NEXT Program), and D.K.B. was supported by US NIH grant GM50936.

## Author’s contribution

M.S., D.K.B., and A.S. designed experiments. M.S. performed all experiments and analyzed the data. M.S., D.K.B, and A.S. prepared the manuscript.

---

**Table S1.**
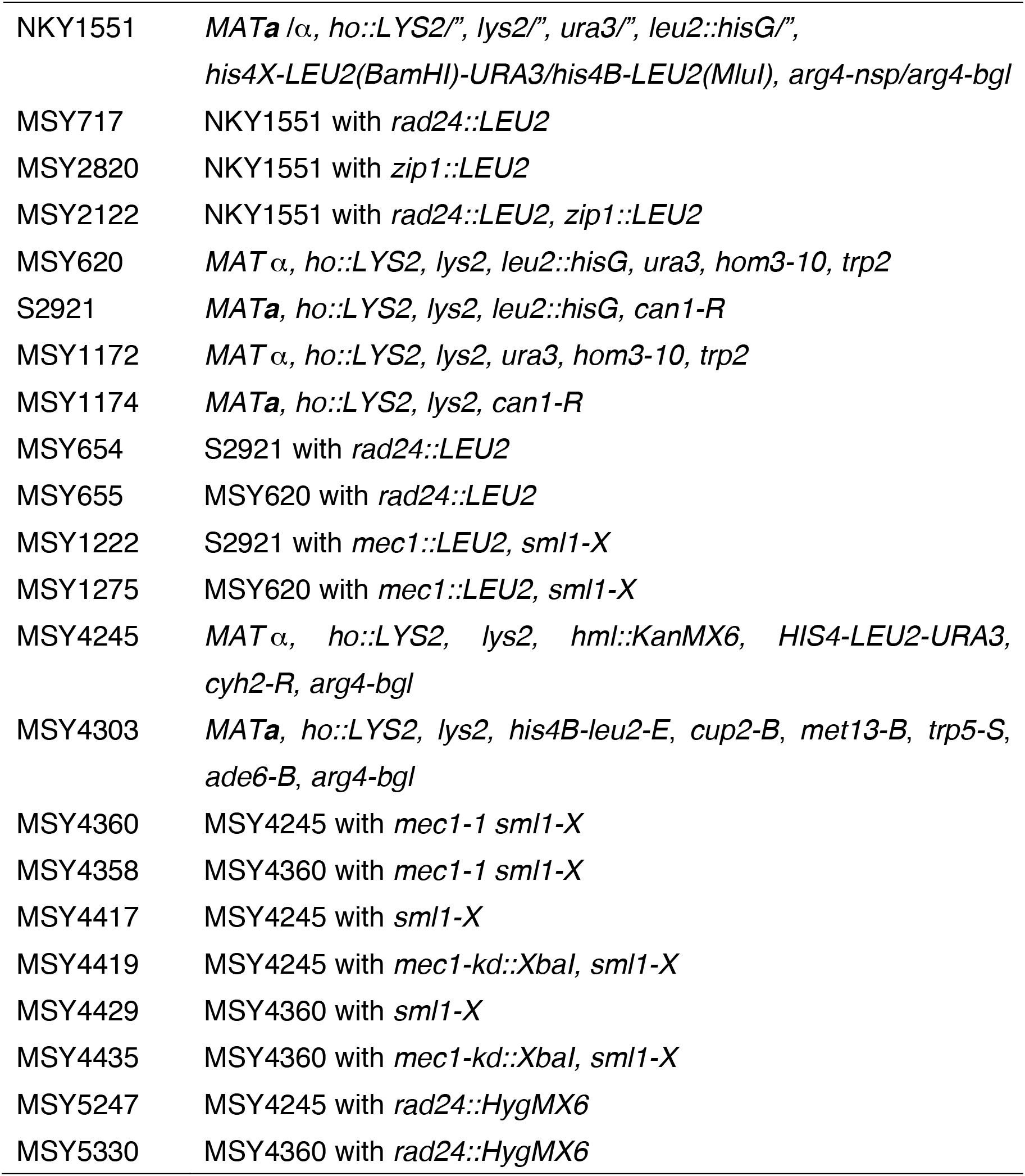
Strain list

**Table S2.**
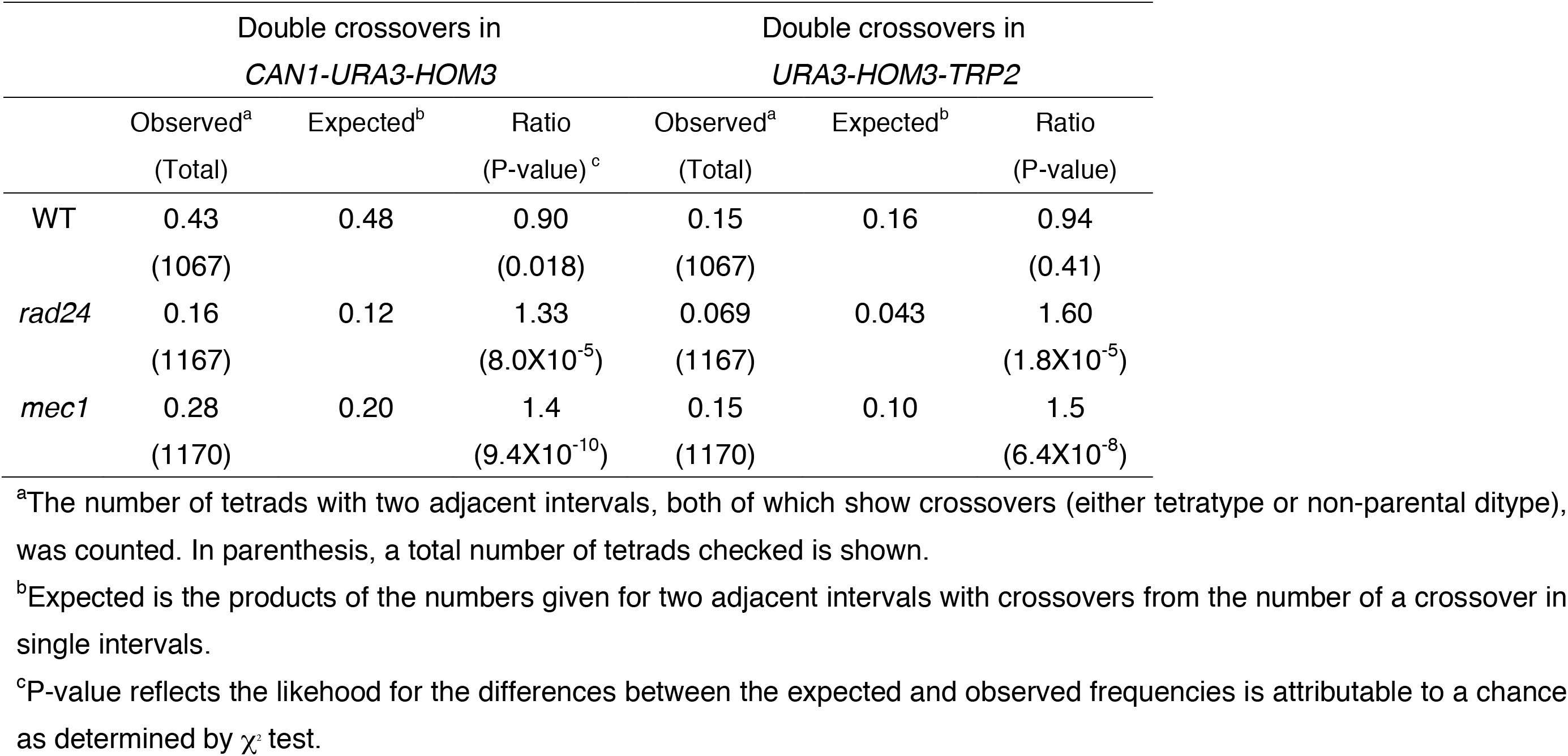
Coincidence of crossovers on chromosome *V*

|  | Double crossovers in<br>*CAN1-URA3-HOM3* |  |  | Double crossovers in<br>*URA3-HOM3-TRP2* |  |  |
| --- | --- | --- | --- | --- | --- | --- |
|  | Observed<sup>a</sup><br>(Total) | Expected<sup>b</sup> | Ratio<br>(P-value)<sup>c</sup> | Observed<sup>a</sup><br>(Total) | Expected<sup>b</sup> | Ratio<br>(P-value) |
| WT | 0.43<br>(1067) | 0.48 | 0.90<br>(0.018) | 0.15<br>(1067) | 0.16 | 0.94<br>(0.41) |
| *rad24* | 0.16<br>(1167) | 0.12 | 1.33<br>(8.0X10<sup>-5</sup>) | 0.069<br>(1167) | 0.043 | 1.60<br>(1.8X10<sup>-5</sup>) |
| *mec1* | 0.28<br>(1170) | 0.20 | 1.4<br>(9.4X10<sup>-10</sup>) | 0.15<br>(1170) | 0.10 | 1.5<br>(6.4X10<sup>-8</sup>) |

<sup>a</sup>The number of tetrads with two adjacent intervals, both of which show crossovers (either tetratype or non-parental ditype), was counted. In parenthesis, a total number of tetrads checked is shown.
<sup>b</sup>Expected is the products of the numbers given for two adjacent intervals with crossovers from the number of a crossover in single intervals.
<sup>c</sup>P-value reflects the likelihood for the differences between the expected and observed frequencies is attributable to a chance as determined by $\chi^2$ test.

**Table S3.** Coincidence of crossovers on chromosome *III*

|  | Double crossovers in<br>*HML-URA3-LEU2* |  |  | Double crossovers in<br>*URA3-LEU2-HIS4* |  |  | Double crossovers in<br>*LEU2-HIS4-MAT* |  |  |
| --- | --- | --- | --- | --- | --- | --- | --- | --- | --- |
|  | Observed<sup>a</sup><br>(Total) | Expected<sup>b</sup> | Ratio<br>(P-value) | Observed<sup>a</sup><br>(Total) | Expected<sup>b</sup> | Ratio<br>(P-value) | Observed<sup>a</sup><br>(Total) | Expected<sup>b</sup> | Ratio<br>(P-value) |
| WT | 0.033<br>(1104) | 0.061 | 0.53<br>(1.7X10<sup>-4</sup>) | 0.0091<br>(1104) | 0.013 | 0.7<br>(0.26) | 0.021<br>(1212) | 0.03 | 0.7<br>(0.084) |
| *rad24* | 0.0078<br>(1160) | 0.0078 | 1<br>(1) | 0<br>(1160) | 0.0012 | NA | 0.0078<br>(1319) | 0.0050 | 1.6<br>(0.18) |
| *mec1-1* | 0.011<br>(1061) | 0.014 | 0.79<br>(0.41) | 0.0038<br>(1061) | 0.0034 | 1.1<br>(0.82) | 0.0076<br>(1213) | 0.01 | 0.76<br>(0.42) |
| *mec1-kd* | 0.039<br>(1033) | 0.034 | 1.1<br>(0.38) | 0.012<br>(1033) | 0.0085 | 1.4<br>(0.22) | 0.015<br>(1226) | 0.022 | 0.7<br>(0.13) |

<sup>a</sup>The number of tetrads with two adjacent intervals, both of which show crossovers (either tetratype or non-parental ditype), was counted. In parenthesis, a total number of tetrads checked is shown.
<sup>b</sup>Expected is the products of the numbers given for two adjacent intervals with crossovers from the number of a crossover in single intervals.
<sup>c</sup>P-value reflects the likelihood for the differences between the expected and observed frequencies is attributable to a chance as determined by $\chi^2$ test.

**Table S4.** Coincidence of crossovers on chromosome *VII*

|  | Double crossovers in<br>*CUP2-MET13-CYH2* |  |  | Double crossovers in<br>*MET13-CYH2-TRP5* |  |  | Double crossovers in<br>*CYH2-TRP5-ADE6* |  |  |
| --- | --- | --- | --- | --- | --- | --- | --- | --- | --- |
|  | Observed<sup>a</sup><br>(Total) | Expected<sup>b</sup> | Ratio<br>(P-value) | Observed<sup>a</sup><br>(Total) | Expected<sup>b</sup> | Ratio<br>(P-value) | Observed<sup>a</sup><br>(Total) | Expected<sup>b</sup> | Ratio<br>(P-value) |
| WT | 0.070<br>(1054) | 0.12 | 0.57<br>(3.7X10<sup>-5</sup>) | 0.13<br>(1054) | 0.15 | 0.86<br>(0.094) | 0.45<br>(1054) | 0.47 | 0.95<br>(0.34) |
| *rad24* | 0.099<br>(1092) | 0.087 | 1.1<br>(0.17) | 0.12<br>(1092) | 0.11 | 1.10<br>(0.32) | 0.41<br>(1092) | 0.38 | 1.1<br>(0.11) |
| *mec1-1* | 0.13<br>(932) | 0.14 | 0.93<br>(0.41) | 0.16<br>(932) | 0.16 | 1<br>(1) | 0.47<br>(932) | 0.48 | 0.98<br>(0.66) |
| *mec1-kd* | 0.15<br>(955) | 0.17 | 0.87<br>(0.13) | 0.18<br>(955) | 0.20 | 0.90<br>(0.17) | 0.53<br>(955) | 0.53 | 1<br>(1) |

<sup>a</sup>The number of tetrads with two adjacent intervals, both of which show crossovers (either tetratype or non-parental ditype), was counted. In parenthesis, a total number of tetrads checked is shown.
<sup>b</sup>Expected is the products of the numbers given for two adjacent intervals with crossovers from the number of a crossover in single intervals.
<sup>c</sup>P-value reflects the likelihood for the differences between the expected and observed frequencies is attributable to a chance as determined by $\chi^2$ test.

**Figure S1.**
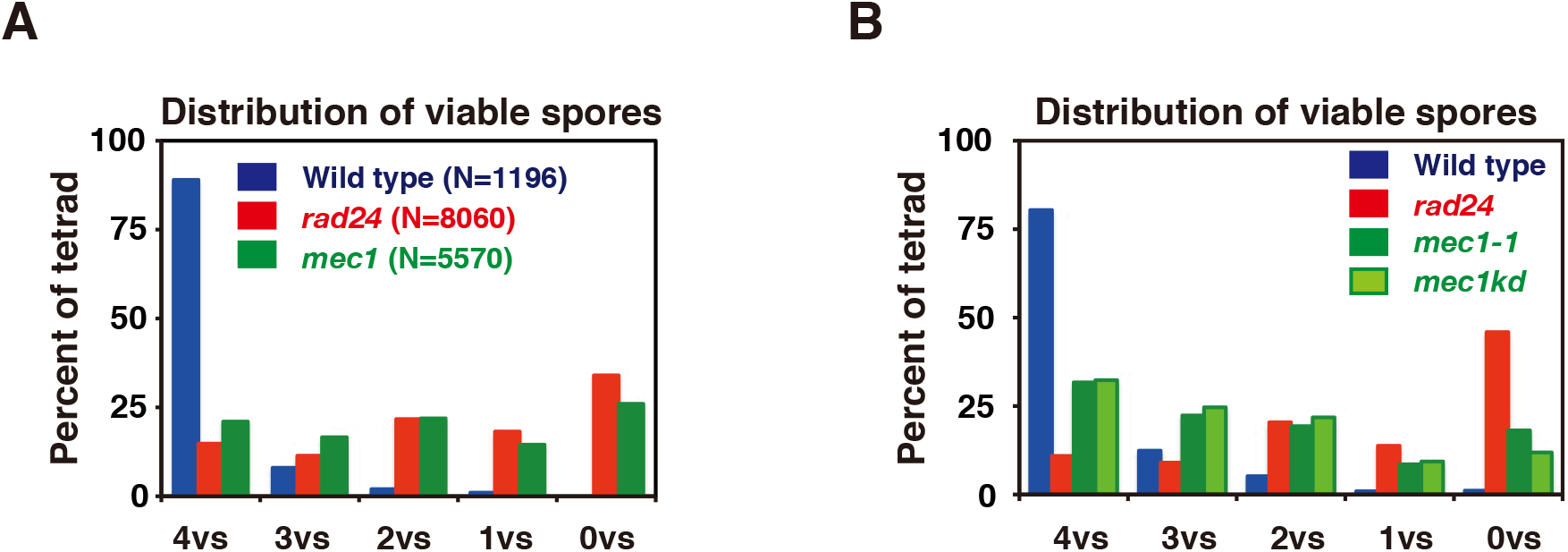
Distribution of viable spores in the *rad24* and *mec1* mutants. A. Distribution of viable spores per tetrad. Numbers of viable spores per tetrad were calculated for wild type (blue bars), the *rad24* (magenta bars) and *mec1 sml1-X* mutants (green bars).
B. Distribution of viable spores per tetrad. Numbers of viable spores per tetrad were calculated for wild type (blue bars), the *rad24* (magenta bars), *mec1-1 sml1-X* mutants (green bars) and *mec1-kd sml1-X* mutants (light green bars).

